# Continuous rearrangement of the postsynaptic gephyrin scaffolding domain: a super-resolution quantified and energetic approach

**DOI:** 10.1101/193698

**Authors:** Pamela C. Rodriguez, Leandro G. Almeida, Antoine Triller

## Abstract

Synaptic function and plasticity requires a delicate balance between overall structural stability and the continuous rearrangement of the components that make up the presynaptic active zone and the postsynaptic density (PSD). Photoactivated localization microscopy (PALM) has provided a detailed view of the nanoscopic structure and organization of some of these synaptic elements. Still lacking, are tools to address the morphing and stability of such complexes at super-resolution. We describe an approach to quantify morphological changes and energetic states of multimolecular assemblies over time. With this method, we studied the scaffold protein gephyrin, which forms postsynaptic clusters that play a key role in the stabilization of receptors at inhibitory synapses. Postsynaptic gephyrin clusters exhibit an internal microstructure composed of nanodomains. We found, that within the PSD, gephyrin molecules continuously undergo spatial reorganization. This dynamic behavior depends on neuronal activity and cytoskeleton integrity. The proposed approach also allowed access to the effective energy responsible for the tenacity of the PSD despite molecular instability.

**Significant statement:** Super-resolution microscopy has become an important tool for the study of biological systems, allowing detailed, nano-scale structural reconstruction, single molecule tracking, particle counting, and interaction studies. However, quantification tools that take full advantage of the information provided by this technology are still lacking. We describe a novel quantification method to obtain information related to the size, directionality, dynamics, and stability of clustered structures from super-resolution microscopy. With this method, we studied the stability of gephyrin clusters, the main inhibitory scaffold protein. We found that gephyrin molecules continuously undergo reorganization based on neuronal activity and changes in the cytoskeleton.

## Introduction

Conventional live-cell fluorescence microscopy has provided considerable insight into the motion of multimolecular assemblies of synaptic proteins. The ensemble diffusion of proteins can be characterized through the use of fluorescent organic molecules or fluorescent proteins. Individual proteins can also be tracked using these dyes or non-organic quantum dots (Dahan et al., 2003). More recently, super-resolution techniques such as photoactivated localization microscopy (PALM) (1) and stochastic optical reconstruction microscopy (STORM) (2) have been applied beyond their initial use for structural reconstruction and used to characterize the nanoscopic organization and diffusion properties of individual proteins within a given structure (3–7). Efforts are now focusing on the development of analytical tools to quantitatively analyze PALM datasets. Pair correlation PALM can describe the organization of proteins within the plasma membrane (8). Recently, PALM has been used for counting proteins within given structures (9, 10). Still lacking is a method to quantify the stability, morphing, and interaction between molecules within nanostructures in live systems.

Synaptic transmission relies on the communication between the presynaptic active zone and the postsynaptic density (PSD). Both specialized structures are composed of thousands of proteins, many of which assemble as clusters at or in close proximity to the synaptic membrane. These clusters are relatively stable over time and exhibit continuous micro-movements and activity-dependent structural changes critical for synaptic plasticity and their alteration can lead to a number of neuropsychiatric diseases (11).

At inhibitory synapses, gephyrin is the central scaffold protein and binds glycine and some GABA_A_ receptor subunits at PSDs (12). The interactions between neurotransmitter receptors and postsynaptic scaffold proteins modulate the distribution and diffusion of receptors in the postsynaptic and extrasynaptic sites (13, 14), which in turn impact neurotransmission strength and synaptic plasticity (15). Gephyrin and inhibitory receptor complexes exhibit constant dynamic fluctuations. FRAP experiments have shown that approximately 30-40% of gephyrin molecules in a postsynaptic cluster are recycled within 5 to 30 minutes (14, 16). This means that within the time frame of a few minutes, most of the changes observed are due to internal molecular rearrangements rather than the entry and exit of gephyrin molecules. As a result, the synaptic scaffold can be used to follow the effective stability of the postsynaptic domain.

Previous studies revealed that postsynaptic gephyrin clusters exhibit sub-micrometric movements on a time-scale of seconds to minutes (17, 18). More recently, we have shown that gephyrin clusters are composed of subdomains, some of which move with respect to each other (10). Reconciling their internal dynamics with the movement of individual gephyrin proteins in relation with synaptic stability and plasticity required a novel approach. Hence, we introduce broadness, elongation, morphing, and effective binding energy as a set of parameters that take advantage of the information obtained from PALM. These parameters quantify changes in directionality, size, intermolecular forces, and dynamics within clustered structures. The proposed method was used to study clusters of postsynaptic gephyrin, tagged with the photoconvertible mEos2 protein and expressed in cultured spinal cord neurons using a lentiviral vector. We investigated the dynamic behavior of PSDs under control conditions and after exposure to pharmacological treatments that tune synaptic activity and disrupt cytoskeletal elements. This allowed an analysis of gephyrin protein rearrangement within clusters and the quantification of the effective energy stabilizing the PSD. Our approach revealed previously undetected changes in the morphing of postsynaptic gephyrin clusters and the nanodomains within. The method described herein can be extended towards the study of other synaptic proteins present at the pre and postsynaptic densities.

## Results

### Postsynaptic gephyrin clusters at super-resolution

To distinguish between synaptic and non-synaptic gephyrin clusters, presynaptic boutons of neurons expressing mEos2-gephyrin were loaded with FM4-64 dye. All mEos2-gephyrin clusters apposed to FM4-64 labeled presynaptic boutons were considered to be synaptic and exhibited a mean area of 0.097 ± 0.007 μm^2^ (mean ± SEM) (Fig. 1A-C; see Fig. S1 for a complete size distribution of clusters), which is within the range previously reported using super-resolution imaging (10).

**Fig. 1.**
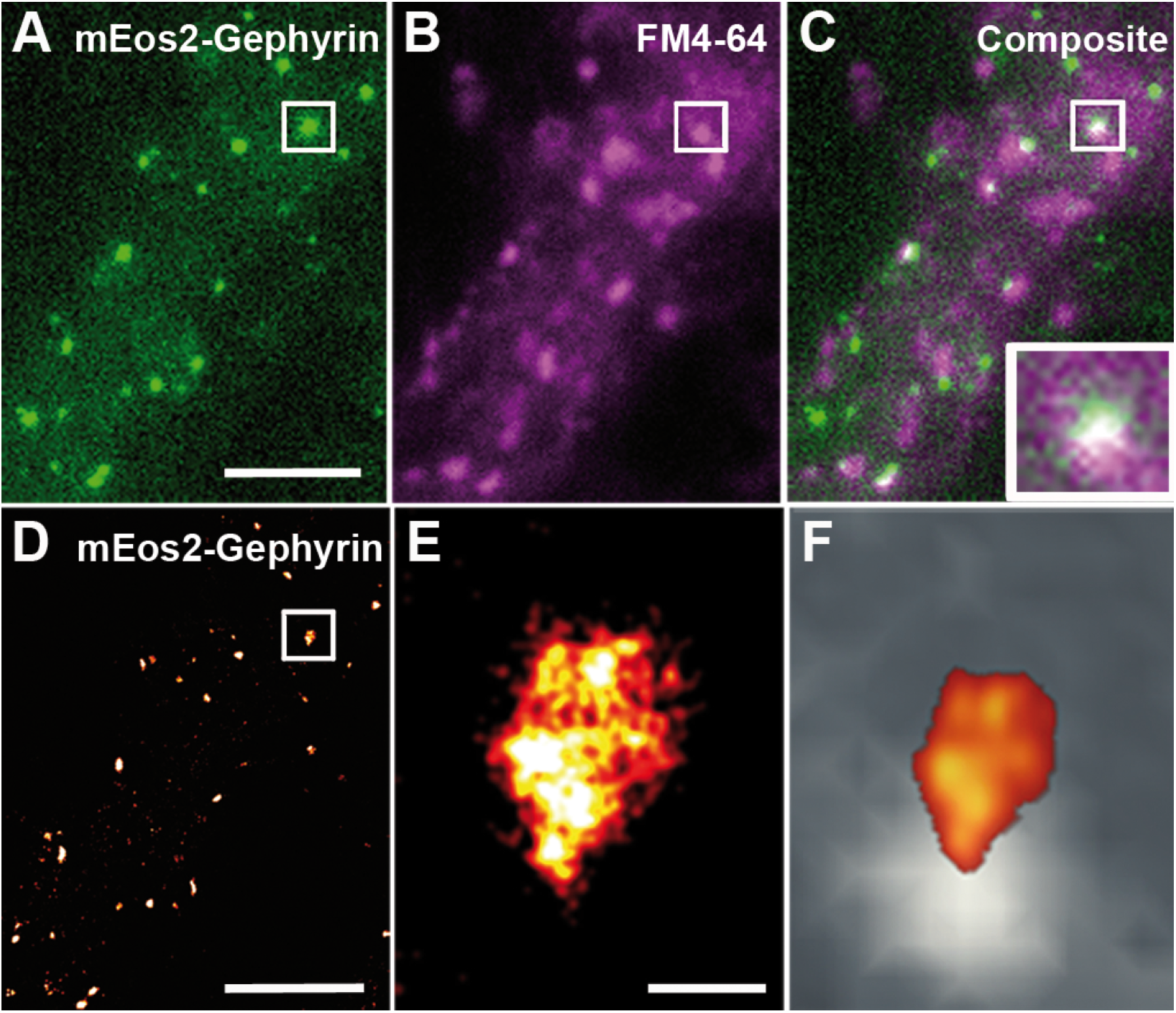
Postsynaptic gephyrin clusters are composed of dynamic sub-domains. (A) Conventional fluorescence microscopy of spinal cord neurons expressing the mEos2-gephyrin protein. Scale bar, 5 µm (B) Presynaptic terminals loaded with FM4-64. (C) Overlay of mEos2-gephyrin and FM4-64 labeling shows a high level of apposition between postsynaptic gephyrin clusters and the presynaptic densities. Higher magnification of the boxed area is shown as an insert. (D) Rendered PALM image of the same field of view as in A. Scale bar, 5 µm (E) Rendered live super-resolution representation of boxed mEos2-gephyrin postsynaptic cluster acquired over a 5.3 minute time period (16000 frames, 20 ms frame rate). Scale bar, 250 nm (F) Smoothed representation of the boxed mEos2-gephyrin postsynaptic cluster (hot-red) superimposed over a smoothed representation of its opposing presynaptic density labeled with FM4-64 (grey scale). (G) Rendered PALM reconstruction of boxed cluster, integrated over 4000 frames with an 80 second temporal resolution. See also Fig. S1.

FM4-64 labeling was used in preliminary experiments. However, because of the photoconvertible nature of the mEos2 fluorescent protein, crosstalk between the FM4-64 and activated mEos2 channels was a concern. In addition, previous studies have shown the existence of functionally silent synapses not labeled by FM4-64 (19), making it a less than ideal marker for our purposes. Therefore, only gephyrin clusters with an area larger than 0.027 μm^2^ under control conditions were used for analysis. On average, cluster area was found to be 0.104 ± 0.005 µm^2^, therefore likely to be synaptic.

A number of postsynaptic gephyrin clusters were composed of sub-domains with varying protein densities (Fig. 1D-F). While whole cluster displacements have been reported in studies using conventional fluorescence microscopy (17, 20), time-dependent PALM on live neurons revealed small displacements of the identified sub-domains with respect to each other (Fig. 1G and 2F). In the following sub-sections, we present a set of analytical tools that make use of PALM data for the study of the internal dynamics and morphing of postsynaptic gephyrin clusters. The use of the analytical method described below can be extended to study the organization, structure, and dynamics of other clustered synaptic proteins, including scaffold assemblies and neurotransmitter receptors.

### Broadness, elongation, morphing, and effective binding energy

The localization precision of individual gephyrin proteins tagged with mEos2 was estimated in fixed samples by measuring the dispersion in the localization of sequential activations belonging to the same fluorophore (σ = 20.91 ± 0.35 nm, mean ± SEM). Using the localization of each detection acquired via PALM, the area of gephyrin clusters and the protein distribution within these clusters were quantitatively described using the second moment of the area with respect to their center of mass (see Eq. 1 in SI Methods) (21). From the second moment of the area, two shape parameters, which describe particular aspects of the distribution of all points in a given area, can be derived: elongation (*E*_*l*_) and broadness (*B*_*r*_) (see Eqs. 2 and 3 in SI Methods) (22). Elongation measures the preferred direction of distribution of gephyrin molecules within a cluster. An elongation value close to zero indicates a preferred linear distribution, while an elongation value close to one indicates a preferred circular distribution. Broadness provides an effective radius of the distribution. An increase in the effective size of a cluster is correlated to an increase in the broadness value, while a decrease in the effective size results in a decreased broadness value. Thus, for a constant number of molecules, the broadness is related to the molecular density.

Using the localization of individual activations, it is thus possible to calculate the broadness and elongation of an assembly of synaptic molecules forming a cluster. Examples of clusters of activations (Fig. 2A-D) that reflect a preferred protein distribution and effective size were quantified using elongation and broadness respectively and plotted in an *E*_*l*_ vs. *B*_*r*_ plane (Fig. 2E). Fig. 2A is representative of the highest population of gephyrin clusters (see Fig. S2 for a complete distribution of cluster *E*_*l*_ and *B*_*r*_). Figure 2F illustrates the time dependent morphological changes of the cluster shown in figure 2C. 3D PALM (see Fig. S3) of postsynaptic gephyrin clusters shows that these 2D observables are congruent with previous findings that describe gephyrin clusters as 2D structures that lie on a plane below the plasma membrane (10, 23).

**Fig. 2.**
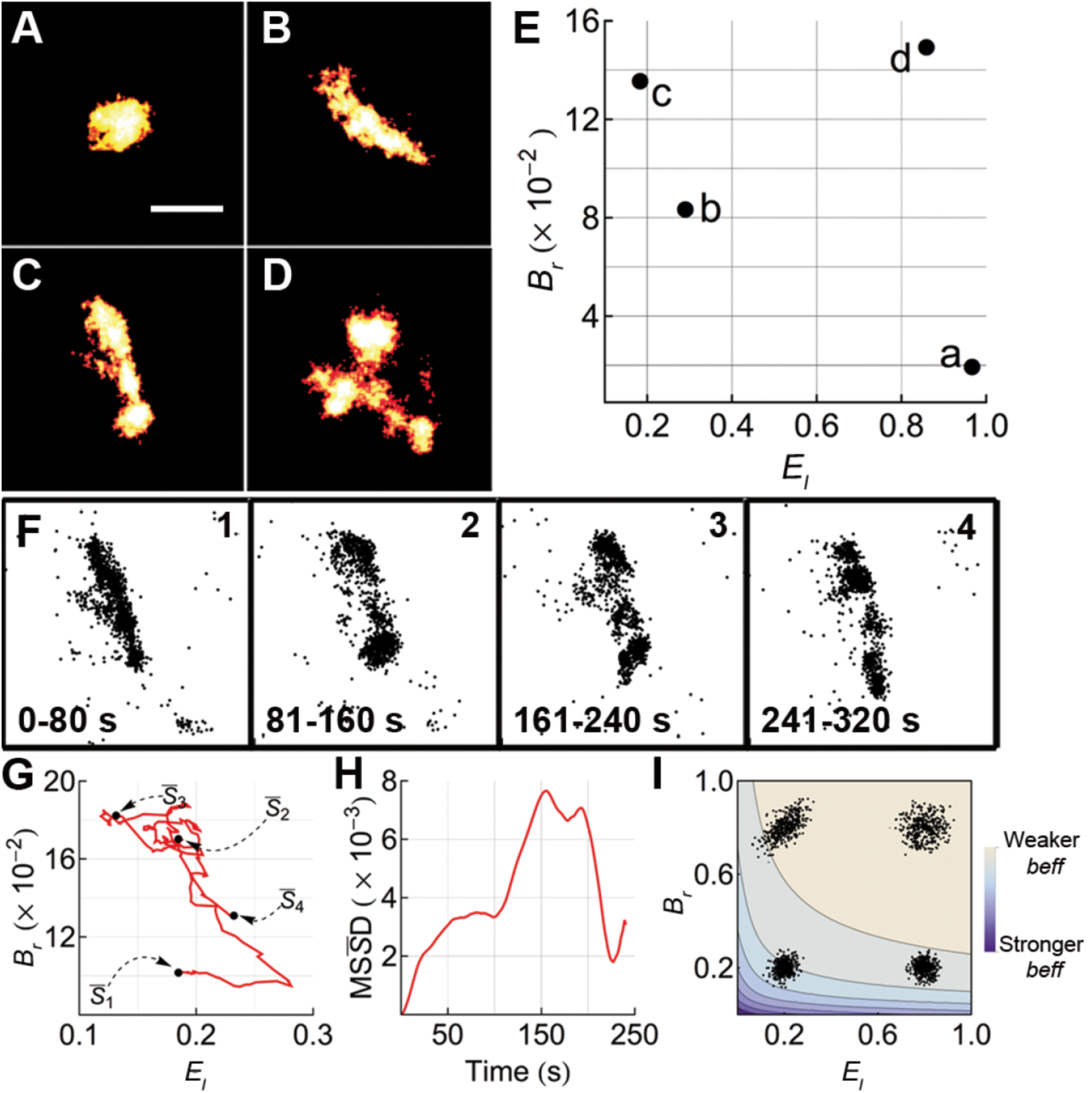
Shape parameters and morphing of postsynaptic gephyrin clusters. (A-D) Representative examples of gephyrin clusters with distinct shape parameters 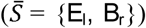. Scale bar, 500 nm (E) Broadness and elongation values of clusters A-D in the 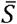 plane. (F) Pointillist representation of cluster C integrated over 4000 frames (80 s) (*F1*: 1-4000 frames, *F2*: 4001-8000 frames, *F3*: 8001-12000 frames, *F4*: 12001-16000 frames). Each point represents a single detection. (G) Localization of the shape parameters 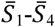 (black dots) of the cluster in panels F1-F4 respectively. The displacement of these shape parameters in the 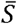 plane over time is shown as a red curve. A sliding window of 4000 frames was used in the calculation of the 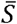 displacement. (H) Mean square shape displacement 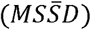 of cluster C. (I) Lines of constant effective binding energy (*b*_*eff*_) on the 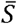 plane. Superimposed are the pointillistic Gaussian distributions with corresponding S parameters. The effective binding energy is weakened as the broadness and/or elongation increase. The shading of the color scale represents the strength of *b*_*eff*_ See also Fig. S2 and S3.

Broadness and elongation can be grouped into a single vector, which we have termed the shape parameter, 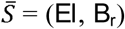. Changes in elongation and broadness over time can be described by the time dependent 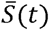 (Fig. 2G). While at a fixed time, the shape parameter provides a snapshot description of cluster shape and size, the time dependent 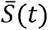 describes the morphological changes that occur within a cluster over time. The relationship between *E*_*l*_ and *B*_*r*_ (shape parameter) quantifies the amount of morphological changes that occur over time. It can be characterized by the mean square displacement of the shape parameter 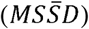 (see Eq. 4 in SI Methods), The shape of the 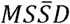 is influenced by variations in elongation and broadness. Figure 2H, illustrates the 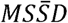 of the cluster shown in figure 2F. From the initial slope of the 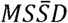 we calculate a morphing parameter 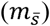, which measures the extent of fluctuations (see Eq. 5 in SI Methods). Low level of variations, represented by low morphing, describe gephyrin clusters that display rather stable shapes over time, while high morphing describes more dynamic clusters and therefore changes in broadness and elongation.

We then used the shape parameter 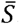 to access cluster stability. Many variables can influence stability, including the molecular density distribution and the affinity between gephyrin and all other PSD components such as receptors and cytoskeletal elements. The relatively long dwell time of gephyrin at synapses allows us to study all these interactions, which can be grouped under an overall effective gephyrin-gephyrin interaction potential termed *b*_*eff*_. The absolute value of this effective potential is a quantitative measure of how strongly cluster components are bound together. A higher value indicates a more stable structure. Fluctuations and a decrease of the absolute value of *b*_*eff*_ may favor morphing and rearrangement of the postsynaptic density.

To access relative changes in *b*_*eff*_ a Gaussian distribution was used since it is a good approximation to model gephyrin distribution within clusters described with a fixed 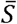. We use the effective potential function linking 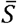 and *b*_*eff*_ (see Eqs. 6 and 10 and the development of the formalism in SI Methods) of this 2D Gaussian distribution as a model for the effective potential (*b*_*eff*_). This *b*_*eff*_ gives a measure of the energetic state (given in units of kB’J) of clusters and its value accounts for the stability of the cluster: a more negative *b*_*eff*_correlates to a more stable cluster. The relation between 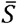 and *b*_*eff*_ constitute isopotential lines and an example is shown in Figure 2I for a particular region of the shape parameter. As a consequence, when broadness and elongation increase, the effective binding between molecules weakens. Note that 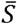 and *b*_*eff*_ values may vary as a function of time. However, the global energetic state of a given cluster remains relatively stable explaining why the structure may remain as a single entity.

### Validation of the physical observables in fixed neurons

When using PALM, a representative image of the structure of interest is reconstructed by grouping multiple frames, each of which contains a small number of mEos2 activations. For dynamic studies, the minimum number of frames required to obtain an accurate representation of the clusters over time must be first determined. To estimate the contribution of the stochastic photo-conversion of the mEos2 tag, independently from molecular movements, data where acquired from fixed spinal cord neurons. When chemically fixed, cluster shape and size should not exhibit changes other than those related to stochastic fluctuations intrinsic to PALM.

To determine the appropriate number of frames of the sliding window to be used, the broadness and elongation of fixed clusters were calculated using windows of varying sizes (1000-15000 frames, 20 ms frame rate). The mean square shape displacement 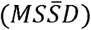 in the *E*_*l*_ vs. *B*_*r*_ plane was then plotted (Fig. 3A). The slope of the 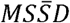 decreased as the size of the sliding window increased due to smaller fluctuations in reconstructing a fixed object. The curve also plateaued for smaller windows, implying that the fluctuations were occurring around a stable shape parameter 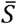. The morphing parameter 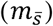 was then calculated while varying the size of the sliding window (Fig. 3B). As expected, the morphing decreased as the number of frames in the sliding window increased. The shape of the morphing vs. sliding window curve (Fig. 3B) is composed of two different scales: an exponential decay between 1000 and 8000 frames, which is followed by a linear decay. The linear portion of the curve indicates that the amount of morphing is stabilized and that the remaining fluctuations are likely due to the spatial stochasticity of mEos2 activations. These results indicate that a minimum of 8000-9000 frames is needed to reconstruct postsynaptic gephyrin clusters. Thus, we used a window of 9000 frames (3 min).

**Fig. 3.**
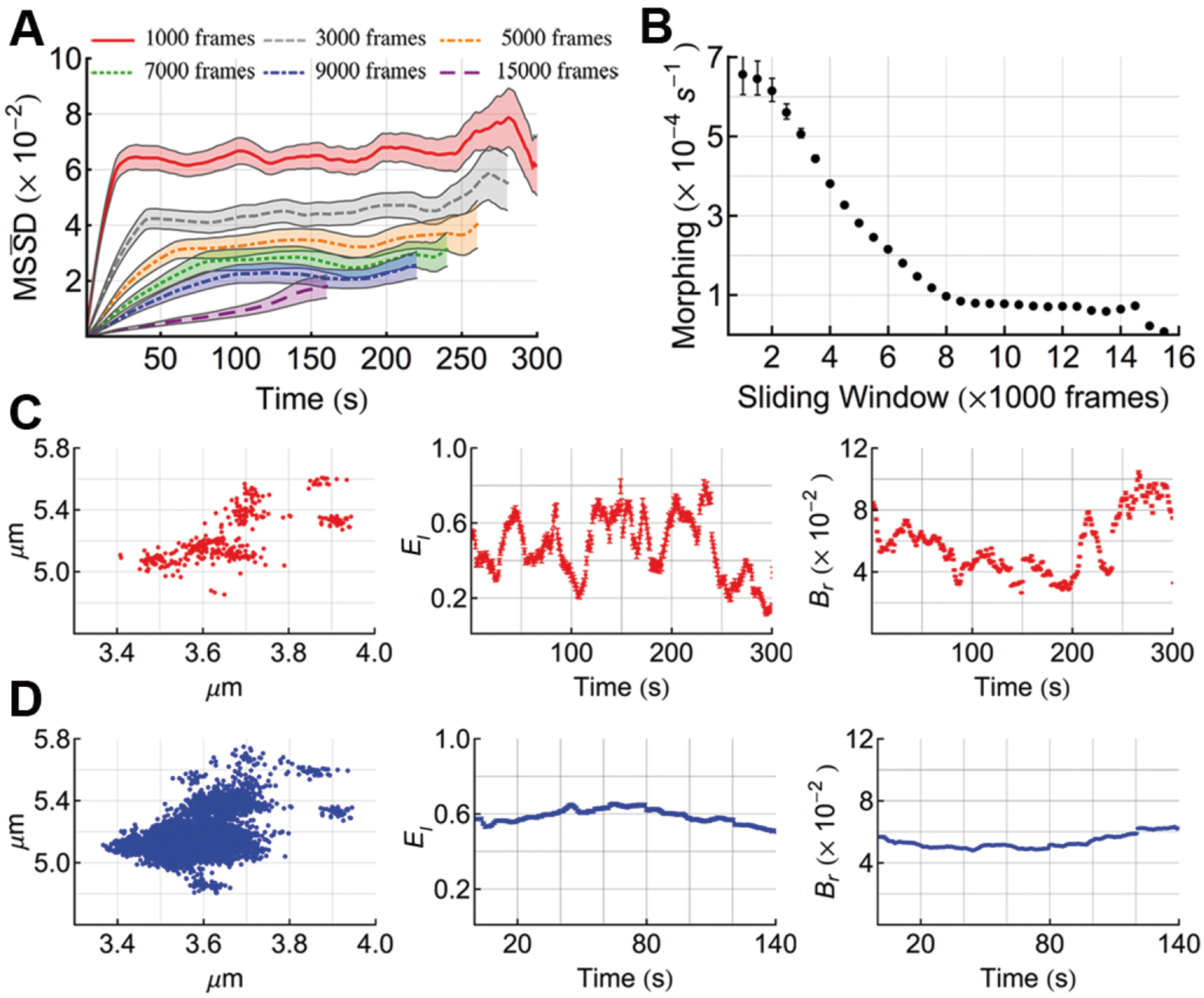
Morphing of postsynaptic gephyrin clusters in fixed spinal cord neurons. (A) 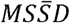 of gephyrin clusters in fixed spinal cord neurons, calculated using sliding windows with increasing number of frames (1000-15000) (B) Morphing parameter of fixed samples with sliding windows of increasing number of frames. The morphing parameter is calculated from the initial slope of the 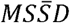 curves. (C) Activations within the first 1000 frames (20 s) of a representative postsynaptic gephyrin cluster (left panel). The elongation (center panel) and broadness (right panel) of the cluster was calculated over time using a sliding window of 1000 frames. (D) Activations within the first 9000 frames (3 min) of the same representative postsynaptic gephyrin cluster as that in C (left panel). The elongation (center panel) and broadness (right panel) of the cluster was calculated over time using a sliding window of 9000 frames. See also Fig. S4 and S5.

The effect of sliding window size and stochasticity are illustrated by following the fluctuations of the shape parameters (*E*_*l*_ and *B*_*r*_) of a representative cluster as a function of time using a sliding window of 1000 frames (20 s, Fig. 3C) and 9000 frames (Fig. 3D). The fluctuations are significantly reduced when using the latter.

The 405 nm laser was set such that the level of activations remained stable over time (Fig. S4 and S5). As a result, the sliding window used for analysis can be set based on number of frames or number of activations. The former was used for our studies.

### Cytoskeletal disruption and alteration of synaptic activity affect gephyrin size, shape, and morphing in live neurons

Alteration of synaptic activity, as well as the disruption of cytoskeletal elements are known to affect the size, shape, dynamics, and lateral movements of postsynaptic gephyrin clusters (17, 24) reflecting molecular reorganization. Therefore, changes in area, elongation, broadness, and morphing of mEos2-gephyrin clusters were compared under control conditions and when treated with nocodazole, latrunculin, 4-AP, or BAPTA-AM. Nocodazole and latrunculin interfere with microtubule and actin polymerization, respectively. The potassium channel blocker, 4-AP, increases synaptic activity by enhancing axonal firing, while BAPTA-AM acts as an intracellular calcium chelator.

First, the surface area of synaptic clusters was determined by using the 90% quantile region of the clusters at 9000 divisions (Fig. 4A). Compared to control, which exhibited a surface area of 0.104 ±0.005 µm^2^ (mean ± SEM), the surface area of gephyrin clusters in fixed neurons (0.101 ± 0.007 µm^2^) was not significantly different. Disruption of microtubules with nocodazole slightly increased their surface area (0.109 ± 0.006 µm^2^), while disruption of actin with latrunculin significantly decreased the surface area (0.090 ± 0.003 µm^2^) compared to control. 4-AP and BAPTA-AM application resulted in opposite effects on cluster surface area, with a significant increase (0.129 ± 0.007 µm^2^) and decrease (0.094 ± 0.009 µm^2^) respectively. These results are in accordance with previously published data (24).

**Fig. 4.**
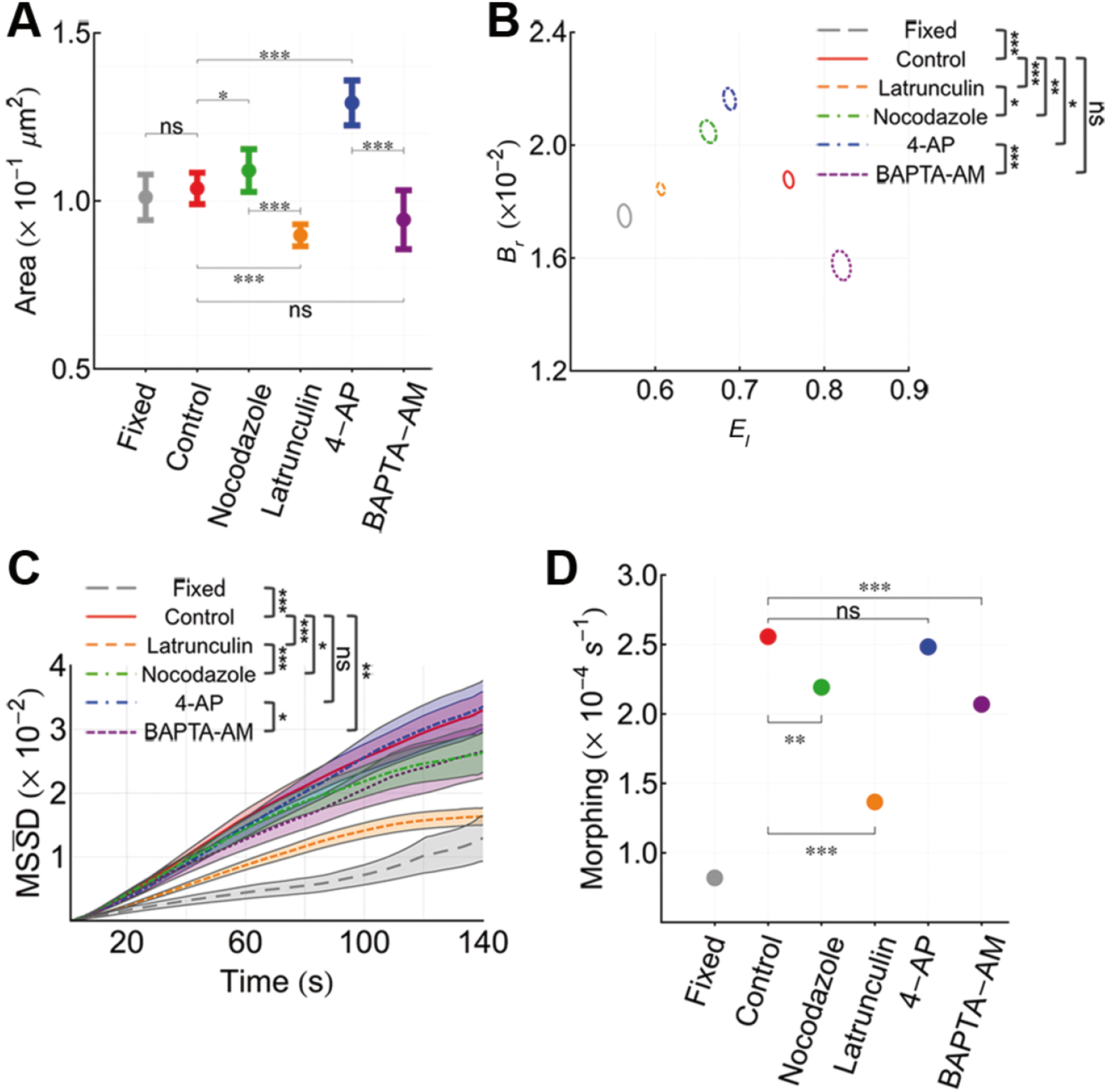
Effect of synaptic activity and disruption of the cytoskeleton on the dynamics of postsynaptic gephyrin clusters in live spinal cord neurons. (A) Mean (95% C.L.) area of gephyrin clusters after aldehyde fixation (n=79 clusters from 11 cells and 2 cultures), under live control (n=331 clusters from 30 cells and 8 cultures) conditions, and when live neurons were treated with nocodazole (10µM, 45 min; n=200 clusters from 20 cells and 3 cultures), latrunculin (3µM, 45 min; n=352 clusters from 26 cells and 3 cultures), 4-AP (50µM, 15 min; n=210 clusters from 17 cells and 4 cultures), and BAPTA-AM (30 μM, 30 min; n=98 clusters from 17 cells and 4 cultures). Significance was determined by One-way ANOVA (F=10.3 at ***p<0.001 with n=1271) with relative significances determined by the Tukey–Kramer method (B) Broadness and elongation of fixed, control, and treated clusters with their corresponding 99% C.L. regions. Significance was determined by MANOVA using Pillai’s trace (F=1192 at ***p<0.0001 with n=1271). Relative significances included Bonferroni corrections. (C) Mean (90% C.L.) 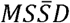 of fixed, control, and treated clusters. Significances were determined by the Mann-Whitney test with Bonferroni corrections. (D) Mean (± SEM) morphing of fixed, control, and treated clusters. Error bars are within the marker size. Significances were determined by Tukey-Kramer method. Fixed clusters were significantly different (***p<0.001) compared to all other measurements. For all legends: *p<0.05, **p<0.01, ***p<0.001

Next, broadness and elongation were computed and plotted in an *E*_*l*_ vs. *B*_*r*_ plane (Fig. 4B). Postsynaptic gephyrin clusters exhibited a distinct morphology under control conditions and following treatment with the same drugs as described above. Fixed neurons exhibited a lower elongation and broadness (0.56 and 0.017, {*E*_*l*_, *B*_*r*_}) compared to control (0.76, 0.019). Treatment with latrunculin resulted in a decrease in both elongation and broadness (0.61, 0.018). Nocodazole led to a decrease in elongation but an increase in broadness (0.66, 0.020). Increasing synaptic activity with 4-AP decreased elongation and increased broadness (0.69, 0.022). Opposite effects were observed when neurons were incubated with BAPTA-AM, leading to an increase in elongation and a decrease in broadness (0.82, 0.016) as compared to control.

A quantitative estimate of cluster morphing can then be obtained by determining the broadness and elongation of clusters over time, done by measuring the 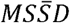 in the *E*_*l*_ vs. *B*_*r*_ plane (Fig. 4C). The morphing of clusters under varying conditions was calculated from the slope of these 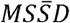 curves (Fig. 4D). Fixed neurons exhibited the lowest level of morphing (0.819 x 10^−4^ ± 0.016 x 10^−4^ s^−1^), consistent with the expected cross-linking of proteins after aldehyde fixation. In live neurons, postsynaptic gephyrin clusters exhibited pharmacological treatment-dependent internal reorganization and dynamics. Compared to control (2.556 x 10^−4^ ± 0.047 x 10^−4^ s^−1^), a significant decrease in morphing was observed when the neurons were treated with nocodazole (2.192 x 10^−4^ ± 0.018 x 10^−4^ s^−1^), latrunculin (1.366 x 10^−4^ ± 0.010 x 10^−4^ s^−1^), and BAPTA-AM (2.069 x 10^−4^ ± 0.009 x 10^−4^ s^−1^), while no significant difference was observed following 4-AP treatment (2.483 x 10^−4^ ± 0.018 x 10^−4^ s^−1^).

Gephyrin molecules within clusters recycle slowly (14, 16), suggesting that the observed changes result from internal molecular rearrangements. Thus, the synaptic scaffold can be used to monitor the effective stability of the postsynaptic region. To approach the effective stability, *E*_*l*_ and *B*_*r*_ were related to the effective binding energy (*b*_*eff*_).

### Accessing the effective energy of protein interactions

The shape parameter region (Fig. 4B) can be overlaid with lines of constant effective binding energy (determined using Eq. 10 in SI Methods) (Fig. 5A). This enables relating the differences in shape parameters to the effective potential between gephyrin proteins. As the name indicates, gephyrin effective energy combines multiple effects into a single potential. That is, it takes into account gephyrin-gephyrin interactions as well as all other gephyrin interacting partners, including receptors, cytoskeletal elements, and other PSD components. Figure 5B provides illustrative examples of the displacement in the *E*_*l*_ vs. *B*_*r*_ plane of three clusters (control (red), latrunculin (orange), and BAPTA-AM (purple)), over the effective binding energy (*b*_*eff*_) isolines. The buffering of calcium with BAPTA-AM resulted in a higher disruption of cluster state, as observed from the shape displacement encompassing multiple *b*_*eff*_ isolines. In contrast, the shape displacement was within fewer or a single *b*_*eff*_ isoline in control and latrunculin treated neurons.

**Fig. 5.**
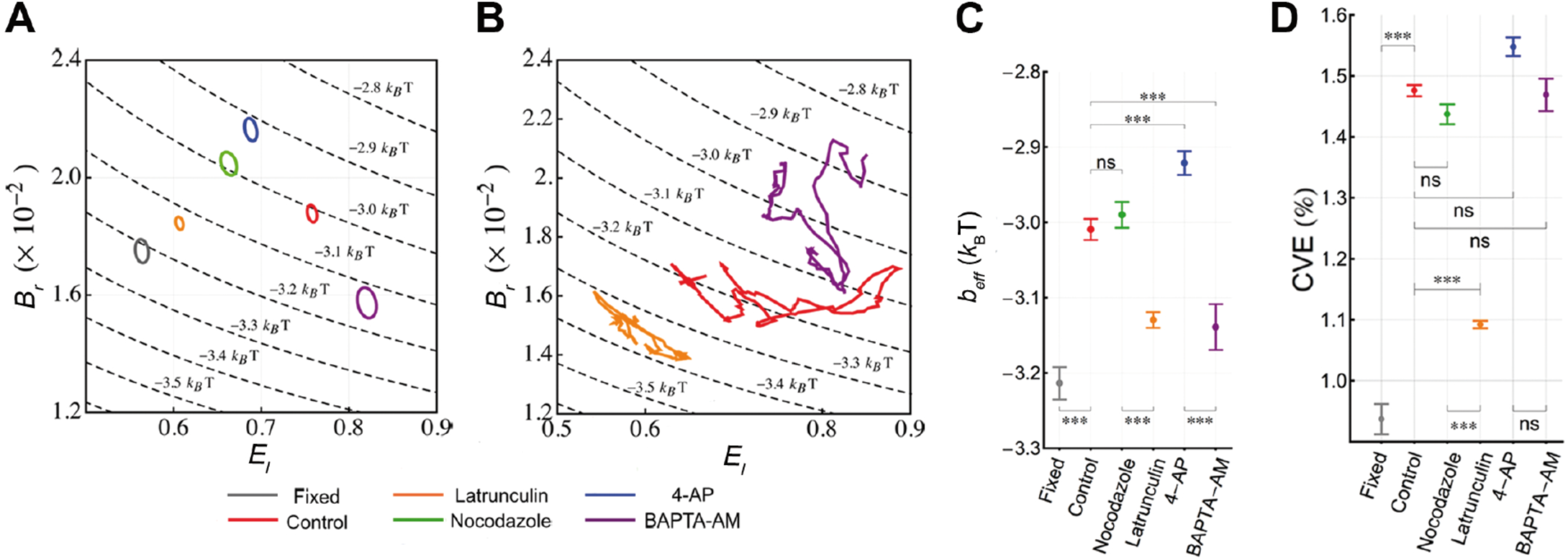
Effective binding energy (bff) of postsynaptic gephyrin. (A) Shape parameter region probed in fixed neurons and live neurons under control and treated conditions overlaid with lines of constant *b*_*eff*_. (B) 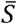 displacement of a control gephyrin cluster (red) and clusters treated with BAPTA-AM (purple) and latrunculin (orange) with respect to the constant *b*_*eff*_. isolines. (C) The inferred mean (99% C.L.) *b*_*eff*_. of fixed, control, and treated gephyrin clusters from the 99% C.L. regions of mean shape parameters in A. Relative significances were determined by the Tukey-Kramer method. (D) Mean (99% C.L.) coefficient of variation of fixed, control, and treated neurons. Significances were determined by One-way ANOVA test (F=12.86 at ***p<0.001 with n=1271). Relative significances determined using the Tukey-Kramer method. For all legends: *p<0.05, **p<0.01, ***p<0.001

The mean effective binding energies (Fig. 5C) provide a measure of how tightly a clustered system is bound: a more negative *b*_*eff*_ correlates to a more tightly bound and therefore more stable cluster. Fixed neurons, exhibited the strongest effective binding energy (-3.214 ± 0.013 k_B_T), consistent with chemical fixation. The binding energy under nocodazole treatment (−2.990 ± 0.010 k_B_T) was not significantly different to that of control (−3.009 ± 0.008 k_B_T). In contrast *b*_*eff*_ increased following 4-AP treatment (−2.921 ± 0.009 k_B_T), indicating that the binding between proteins is weaker and less stable, therefore making the clusters easier to disrupt. This is consistent with their increased surface area (Fig. 4A). In comparison, treatment with latrunculin and BAPTA-AM led to clusters with a significant decreased *b*_*eff*_ (−3.129 ± 0.006 k_B_T and −3.139 ± 0.018 k_B_T respectively), making them more stable, harder to disrupt, and more compact (Fig. 4A).

For dynamic studies, changes in *b*_*eff*_ over time can be followed using a coefficient of variation (*CVE*) (Fig. 5D), which provides a measure of the variance of the effective binding between gephyrin molecules within individual clusters. Fixed neurons exhibited the lowest coefficient of variation. For control and treated neurons, we found that the variations were within an order of 1%, implying that the cluster shape parameter movements are confined within specific *b*_*eff*_ isolines. This type of confined movement is illustrated by the examples (Fig. 5B) of control (red) and latrunculin (orange) treated clusters, while BAPTA-AM (purple) treatment illustrates movement with higher fluctuations. While both *CVE* and 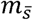 provide a measure of how much the shape parameters are changing, only *CVE* provides information regarding the movement along the *b*_*eff*_ isolines. Therefore,*b*_*eff*_ reflects the stability of the structure and its fluctuation (CVE) provides a measure of the capacity to shift from one stable state to another.

## Discussion

Broadness, elongation, morphing, and effective binding energy are quantitative tools developed to characterize the structural and molecular dynamics of and within clustered structures. The application of the method is demonstrated here for the study of postsynaptic gephyrin clusters imaged in live neurons by super-resolution microscopy.

The PSD can be maintained for a long time by far exceeding the dwell time of individual molecules in the structure (25). The continuous binding and unbinding of molecules is at the origin of the morphing activity, which is affected by synaptic activity or by disruption of the cytoskeleton elements. Optical techniques such as FRAP and FLIP have been used to measure the mobility of molecules, including that of gephyrin and other scaffold proteins at synapses (14, 16, 26, 27). Although valuable, these tools measure population dynamics and provide no information regarding the displacement and micro-organization of individual molecules.

Since its development, PALM has been used to unravel the micro-organization of proteins and other biological molecules. Besides gephyrin, PALM has revealed the nanodomain organization of other PSD proteins including AMPA receptors (4) and PSD-95 (7). In recent years, more interest has been placed on the development of analytical tools to extract information regarding the number (8–10), diffusion (3–5), and interaction (28, 29) of molecules. However, none of the analytical methods currently available have been used to quantify the stability, morphing, and interaction between molecules within small (approximately 0.1µm^2^) clustered structures in live systems.

First, we show that the morphing parameter introduced herein, can be used to determine the minimum number of activations (or frames) needed to accurately reconstruct a super-resolved PALM image in live neurons. This is important since undersampling, together with the stochastic nature of PALM, can lead to artifacts such as large fluctuations of shape and size over short periods of time. Analysis of fixed neurons showed that it was not necessary to image the fluorescently tagged clusters to exhaustion to obtain a representative picture of their structure.

Postsynaptic gephyrin clusters under control and treated conditions, exhibit a distinct shape and size, observed as a localization of 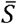 in the *E*_1_ vs. *B*_*r*_ plane (Fig. 4B). Although the clusters did not exhibit drastic changes during the time period of acquisition, the relative position of the molecules within the clusters changed constantly. Alterations of tubulin, actin, or synaptic activity, regulated the morphing of the PSD and internal movements. Thus, despite an apparent stability of the postsynaptic gephyrin clusters, our analytical method, together with PALM, reveals nanoscopic rearrangements. These small and continuous rearrangements may facilitate a change in scaffold protein interactions ultimately accounting for synaptic plasticity (30, 31).

We further related the distinct shape parameters to a dynamical component of the clustering proteins, the effective binding energy. This energy represents gephyrin pair interactions and takes into account all other interacting partners, including receptors, cytoskeletal elements, and other PSD components (28, 29). In this context, changes in effective binding energy were actin-, microtubule-, activity-, and calcium-dependent and regulated clustering. Together with the coefficient of variation, which quantifies the energy fluctuation, we can obtain a measure of stability in the postsynaptic density. A lower coefficient of variation, such as that seen after actin fiber disruption, indicates that the binding between molecules is steadier, making the structure less likely to morph. It is hypothesized that gephyrin forms a hexagonal lattice composed of strongly interacting trimers, which can themselves dimerize with lower affinity (32–34). Therefore, the effective binding energy variation is likely to result from fluctuations of the dimeric interaction.

An important advantage of the proposed method of analysis, used in conjunction with super-resolution imaging, is the ability to obtain a more in-depth understanding of the system under investigation. Using our approach, it is now possible to detect small yet significant differences that would not be discerned with conventional fluorescence microscopy. While area, effective binding energy, and coefficient of variation appear to be correlated, our results suggest this is not always the case. Treatment with BAPTA-AM, for example, did not significantly affect cluster area compared to control. However, a significant reduction in effective binding energy was observed after calcium buffering. Studies performed using wide-field fluorescence microscopy suggested that actin fibers and microtubules have opposite effects on gephyrin dynamics, molecular density, and size (17, 24). Our method quantifies the amount of micro-movements that occur within gephyrin clusters as well as their stability. Calcium influx was shown to stabilize spines by suppressing actin dynamics (35). Our results now show that both actin depolymerization by latrunculin and calcium chelation by BATPA - AM result in decreased morphing and effective binding energy of gephyrin clusters.

Proteins that make up the postsynaptic density of excitatory synapses have also been shown to exhibit dynamic properties with similar recycling times as those observed in inhibitory synapses. PSD-95, for example, exhibits a 20% turnover within 5 minutes (36). Therefore, this analysis has immediate application in the neuroscience field for both inhibitory and excitatory synapses. Its use will provide a quantitative approach to study the morphology, the time dependent morphing, and the energetic state of synaptic clusters in relation with synaptic stability and plasticity.

While the parameters described herein can be used to study various types of clustered structures, they are not comprehensive. Investigating higher order moments could provide additional information. Furthermore, these parameters need to be adapted and limitations intrinsic to the studied molecules need to be taken into account. More precisely, in order to approach the dynamic properties of a cluster of proteins, the acquisition time has to be set depending on the characteristic time of recycling of the structure and of its components. This will reinforce the notion that the changes observed result from structural dynamics and not from the entry and exit of the molecules into and out of the studied structure. In the case of gephyrin, for example, the recording time of 7 min with an acquisition rate of 20 ms is well below its characteristic time, which is of approximately 30 min.

The recording time is dependent on the molecular counts and must be increased or decreased accordingly. A structure with high molecular counts needs to be imaged for a longer period of time for an accurate and reliable reconstruction. The proposed analysis can be used for structures of any shape and size; the only limitation is the optical resolution and the field of view of the microscope. However, there must be at least two molecules within each sliding window in order to define the shape parameters. This is the lowest theoretical limit but practically, the actual number depends on the number of occurrences in a given sliding window. Importantly, the molecular counts must remain constant over sliding windows. The model used for the quantification of the effective energy can be used for any structure that can be described by a Gaussian distribution.

This quantification approach can be applied to planar, globular, and filamentous structures, and in certain cases, 3-dimentional (3D) imaging should be considered. Implementing adaptive optics, it is possible to obtain an axial resolution of 40 nm (37). For planar structures such as the membrane-associated gephyrin, which is known to lie on a plane below the plasma membrane (10, 23), the 3D aspect does not have a significant influence on the results when the same element is studied over time (see Fig. S3). For globular structures, 3D imaging is critical since the quantitative method depends on the distance between points, which can be large in 3D structures, placing more importance on the angle of projection and its change over time. If the structure of interest is filamentous, 3D imaging should also be taken into consideration since the filament can span in various planes and have tortuous movements.

In conclusion, we have demonstrated the application of broadness, elongation, morphing, and effective energy for the analysis of postsynaptic gephyrin clusters in live neurons. This allows the quantification of internal changes and stability. The stability can now be quantified in *k*_B_T and provides a means to access the effective energy contributing to the tenacity of the structure over time. This means that depending on the physiological condition, a given synapse may be more or less prone to shift from one status of quasi-equilibrium to another (38), thus defining the concept of meta-stability.

## Methods

### Drugs and Reagents

Unless otherwise noted, all chemicals were purchased from Sigma-Aldrich (St. Louis, MO) or Life Technologies/Molecular Probes (Carlsbad, CA).

### Lentivirus

A lentivirus encoding mEos2-Gephyrin was produced by cotransfecting the lentiviral backbone plasmid (FUGW) encoding the mEos2-Gephyrin construct (5 µg) along with the pMD2.G envelope (5 µg) and pCMVR8.74 packaging (7.5 µg) plasmids (Addgene, Cambridge, MA) into HEK293T cells using lipofectamine 2000 (60 µl). Transfection was performed in 10 cm plates once the cells reached 80% confluence. Supernatant containing lentivirus was collected 48 h after transfection, filtered through a 0.45 µm-pore-size filter, aliquoted, and stored at –80 °C.

### Cell culture and infection

All experiments were performed on dissociated spinal cord neuron cultures prepared from Sprague-Dawley rats (at E14). Experiments were carried out in accordance with the European Communities Council Directive 2010/63EU of 22 September 2010 on the protection of animals used for scientific purpose and our protocols were approved by the Charles Darwin committee in Animal experiment (Ce5/2012/018). Neurons were plated at a density of 6.3 x 10^4^ cells/cm^2^ on 18 mm coverslips pre-coated with 70 µg/ml poly-D,L-ornithine and 5% fetal calf serum. Cultures were maintained in neurobasal medium containing B-27, 2 mM glutamine, 5 U/ml penicillin, and 5 µg/ml streptomycin at 37°C and 5% CO_2_. Neurons were infected at 7 days in vitro (DIV) with a recombinant lentiviral vector expressing the mEos2-gephyrin construct.

### PALM imaging

Photoactivated localization microscopy (PALM) was performed at DIV 14-17 on live neurons with no treatment (control), live neurons treated with 4-AP (50µM, 15 min), BAPTA-AM (30µM, 30 min), nocodazole (10µM, 45 min), or latrunculin (3µM, 45 min), and neurons fixed with 4% paraformaldehyde and 1% sucrose (10 min). Live imaging was performed at 35°C in imaging medium (minimum essential medium with no phenol red, 33 mM glucose, 20 mM HEPES, 2 mM glutamine, 1 mM sodium pyruvate, and B-27). All experiments were performed on an inverted Nikon Eclipse Ti microscope with a 100x oil-immersion objective (N.A. 1.49), an additional 1.5x lens, and an Andor iXon EMCCD camera. Super-resolution movies were acquired at 20 ms frame rate under continuous illumination with activation (405 nm) and excitation (561 nm) lasers for a total of 20000 frames (6.7 minutes). Activation density was kept steady by manually increasing the activation laser intensity over time. Conventional fluorescence imaging was performed with a mercury lamp and filter sets for the detection of preconverted mEos2 (excitation 485/20, emission 525/30). The z-position was maintained during acquisition by a Nikon perfect focus system.

### 3D PALM imaging

3D PALM was performed as described previously using adaptive optics (AO) to induce astigmatism to the PSF of single molecule detections (10, 37). The imaging set-up was as described above, with the addition of a MicAO system (Imagine Optic) in the emission pathway. Calibration curves were taken at the beginning of each experiment using 100 nm TetraSpeck beads, imaged using a nanopositioning piezo stage over a range of 1 µm with a 0.025 µm step size.

Please refer to SI Text for methodology regarding single molecule detection, cluster selection, and data analysis and statistics.

### Author contributions

Conceptualization, P.C.R., L.G.A., and A.T.; Methodology, P.C.R. and L.G.A.; Investigation, P.C.R.; Software, L.G.A.; Formal Analysis, P.C.R. and L.G.A.; Writing – Original Draft, P.C.R. and L.G.A.; Writing – Review and Editing, P.C.R., L.G.A., and A.T.

## ACKNOWLEDGMENTS

We thank S. Edelstein, M. Dahan, and K. Sekimoto for proofreading and constructive criticism of the manuscript and M. Renner for technical support. This work was supported by ERC Plastinhib and ANR Synaptune and the Program `Investissements d’Avenir’: ANR-10-LABX-54 MemoLife. P.C.R. was supported by a Marie Curie International Incoming Fellowship within the 7th European Community Framework Programme and L.G.A. was funded by the European Union Seventh Framework Programme (FP7/2007-2013) under grant agreement number 604102 (HBP). The authors declare no competing financial interests or conflicts of interest.

